# Global transcriptome analysis of *Stenotrophomonas maltophilia* in response to growth at human body temperature

**DOI:** 10.1101/2021.01.10.426099

**Authors:** Prashant P. Patil, Sanjeet Kumar, Amandeep Kaur, Samriti Midha, Kanika Bansal, Prabhu B. Patil

**Affiliations:** Bacterial Genomics and Evolution Laboratory, CSIR-Institute of Microbial Technology, Chandigarh, India; Department of Microbiology, School of Medicine, University of Washington, Seattle, WA, USA; Imperial Life Sciences (P) Limited, Gurgaon, Haryana, 122001, India; Institute of Infection, Veterinary and Ecological Sciences, University of Liverpool, Liverpool, United Kingdom

**Keywords:** RNA-Seq, *Stenotrophomonas maltophilia*, thermoregulation, transcriptome

## Abstract

*Stenotrophomonas maltophilia* (Smal) is a typical example of an environmental originated opportunistic human pathogen, which can thrive at different habitats including the human body and can cause a wide range of infections. It must cope with heat stress during transition from the environment to the human body as the physiological temperature of the human body (37 ◻) is higher than environmental niches (22-30 ◻). Interestingly, *S. rhizophila* a phylogenetic neighbour of Smal within genus *Stenotrophomonas* is unable to grow at 37 ◻. Thus, it is crucial to understand how Smal is adapted to human body temperature, which could suggest its evolution as an opportunistic human pathogen. In this study, we have performed comparative transcriptome analysis of *S. maltophilia* grown at 28 ◻ and 37 ◻ as temperature representative for environmental niches and human body respectively. RNA-Seq analysis revealed several interesting findings showing alterations in gene expression levels at 28 ◻ and 37 ◻, which can play an important role during infection. We have observed downregulation of genes involved in cellular motility, energy production and metabolism, replication and repair whereas upregulation of VirB/D4 Type IV secretion system, aerotaxis, cation diffusion facilitator family transporter and LacI family transcriptional regulators at 37 ◻. Microscopy and plate assays corroborated altered expression of genes involved in motility. The results obtained enhance our understanding of the strategies employed by *S. maltophilia* during adaptation towards the human body.

**Impact statement:** *Stenotrophomonas maltophilia* (Smal) is a WHO listed multidrug resistant nosocomial pathogen. Interestingly, *S. maltophilia* species can grow both at 28 ◻ and 37 ◻ unlike its closest taxonomic relative, i.e., *S. rhizophila* and also majority species belonging this genus. Hence this ability to grow at 37 ◻, i.e., human body temperature might have played key role in the unique success and emergence of this species as opportunistic human pathogen. Using transcriptome sequencing, we have identified set of genes which are differentially regulated at 37 ◻ and investigated their evolutionary history. This study has revealed regulation of genes involved in motility, metabolism, energy, replication, transcription, aerotaxis and a type IV secretion system might have a role in successful adaption to a distinct lifestyle. The findings will be helpful in further systematic studies on understanding and management of an emerging human pathogen such as Smal.

## Introduction

Variation in temperature is one of the most crucial stress factors for pathogens of environmental origin during adaptation to the human body, as temperature of the external biosphere is generally 22-30 ◻. There are different molecular mechanisms by which bacteria sense and respond to changes in temperature. Moreover, temperature is one of the critical signals that influences the different bacterial processes. In bacterial pathogens of mammals including *Shigella*, *Yersinia, Pseudomonas* etc., the body temperature of host, i.e. 37 ◻ induces the expression of virulence factors (White-Ziegler, Malhowski et al. 2007, Wurtzel, Yoder-Himes et al. 2012). Temperature is one of the important signals that a mammalian pathogen uses to regulate the virulence trait once it has entered its warm-blooded host (Konkel and Tilly 2000). In contrast, in pathogens of plants and ectothermic hosts such as fish, molluscs and amphibians, virulence gene expression is elevated at the lower temperatures, suggesting a role of temperature in the coordination of bacterial pathogenesis and virulence (Shapiro and Cowen 2012, Lam, Wheeler et al. 2014). Recently discovered RNA thermometers are an interesting tool in bacteria for responding to such external temperature stresses. They are RNA structures formed at the 5′ UTR regions of transcripts specifying regulatory proteins responsible for expression of virulence-associated traits, which blocks translation initiation of genes at non-permissive temperatures (Kortmann and Narberhaus 2012).

Genus *Stenotrophomonas* comprises several species from diverse range of niches such as *S. lactitubi* and *S. indicatrix* from food, *S. bentonitica* and S. *chelatiphaga* from soil etc. (Patil, Midha et al. 2016, Patil, Kumar et al. 2018). *S. maltophilia* (Smal) is a ubiquitous bacterium which has emerged as multi drug-resistant global opportunistic pathogen in immunocompromised patients (Looney, Narita et al. 2009, Brooke 2012, Brooke 2014). Smal is a versatile bacterium, which adapts a wide range of environments and it is the only validated species among *Stenotrophomonas* genus, which causes human and animal-associated infections (Ryan, Monchy et al. 2009, Patil, Kumar et al. 2018). Apart from this detrimental effect, Smal has an extraordinary range of activities such as plant growth promotion, degradation of anthropogenic pollutants and production of biomolecules (Ryan, Monchy et al. 2009, Mukherjee and Roy 2016). Presence of such a wide range of properties makes this bacterium an important biotechnological candidate, but the pathogenic potential of this bacterium limits its use for biotechnological applications (Mukherjee and Roy 2016). The comparison of the Smal with *S. rhizophila*, a non-pathogenic and phylogenetically related species, revealed that *S. rhizophila* lacks crucial virulence factors and heat shock proteins (Alavi, Starcher et al. 2014). *S. rhizophila* is unable to grow at human body temperature, 37 ◻ due to the absence of heat shock genes and upregulation of genes involved in suicidal mechanisms (Alavi, Starcher et al. 2014). Thus, it is essential to understand the adaptation of rapidly emerging multidrug resistance opportunistic pathogen Smal to human body temperature, which is considered as the first step towards transition from environment to the human body.

Advances in high-throughput sequencing approaches will accurately quantify levels of expression of mRNA (RNA-Seq) thus, providing significant advances over microarrays (Croucher and Thomson 2010, Trapnell, Hendrickson et al. 2013, Creecy and Conway 2015). To understand the genetic response, mechanistic basis and factors involved in the successful adaptation of the *S. maltophilia* at human body temperature, we systematically examined the transcriptome during the growth at 28 ◻ and 37 ◻ using RNA-Seq experiments.

## Methods

### Bacterial strain and growth condition

*S. maltophilia* strain MTCC 434^T^, which is isogenic with the ATCC 13637^T^ was used in all experiments. *S. maltophilia* ATCC 13637^T^ was grown in Luria Bertani Miller Broth with shaking at 200 at either 37 ◻ or 28 ◻.

### Total RNA extraction, Quantification and Integrity estimation

*S. maltophilia* ATCC 13637^T^ was grown in 20 ml Luria Bertani Broth, Miller in 100 ml Erlenmeyer flask at 37 ◻ and 28 ◻ under constant agitation at RPM 200 (Supplementary Fig 1). Samples were withdrawn at intervals for optical density monitoring at 600 nm (OD_600_), and cells from both cultures were harvested at mid-log phase (OD_600_ = 0.8 to 1) by centrifugation at 6000 rpm at for 10 min at 4 ◻ and immediately frozen at −80 ◻ or proceeded to the RNA isolation. For isolation of RNA, the pellet was resuspended in the 1 ml of TRIzol (Invitrogen, Carlsbad, CA, USA) and dissolved by vigorous mixing. The supernatant was transferred into a clean tube which contained one volume of 100% ethanol mixed by repeated gentle inversion. The RNA was purified and treated with DNase by using the Direct-zol RNA MiniPrep kit (Zymo Research Corporation, Orange, CA, USA), according to the manufacturer’s recommendation. The purity of isolated total RNA, was determined by using the NanoDrop (Thermo Scientific, Wilmington, DE, USA) and quantified by using Qubit (Invitrogen, Carlsbad, CA, USA). Agilent Bioanalyzer with Agilent RNA 6000 Nano Kit (Agilent Technologies, Palo Alto, CA, USA) was used as per manufacturer’s guidelines to assess the integrity of RNA samples. The RNA samples with RNA Integrity Number (RIN) > 8 were selected for cDNA synthesis and subsequent Illumina library construction (Supplementary Fig 1).

**Figure 1:**
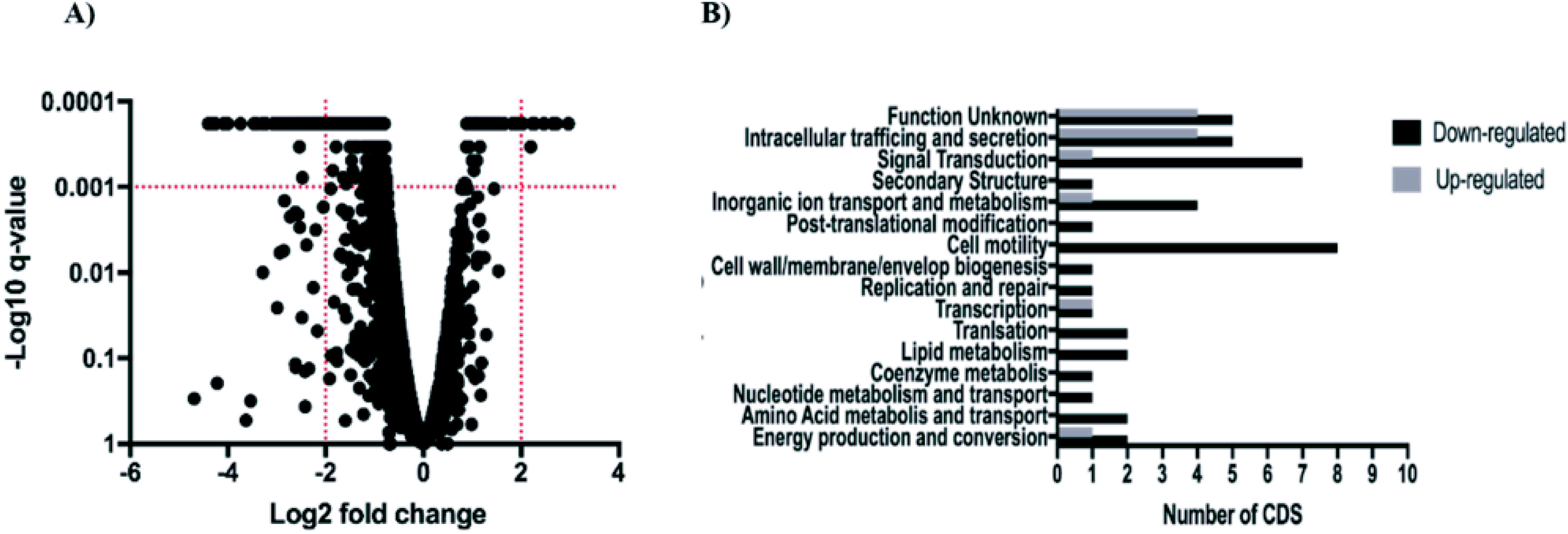
Transcriptional response of*S. maltophilia* ATCC 13637^T^ at 37 °C. A) Volcano plot showing the transcripts that fulfil both fold change (log2 fold) and q-value <0.001 cut-offs. B) COG-based classification of differentially expressed genes of *S. maltophilia* at 37 °C.

### Ribosomal RNA depletion, cDNA library preparation and Illumina sequencing

The ScriptSeq complete kit (Epicentre, Illumina, Madison, WI USA), a combined kit for the ribosomal (rRNA) depletion Ribo-Zero™ Kit (Bacteria) (Epicentre, Illumina, Madison, WI USA) and cDNA library construction kit, ScriptSeq™ v2 RNA-Seq library preparation kit (Epicentre, Illumina, Madison, WI USA) was used for this purpose. Total 5 μg of RNA was used for rRNA depletion by using Ribo-Zero™ (Epicentre, Illumina, Madison, WI USA) kit and purified by using Qiagen-RNeasy miniElute (Qiagen GmbH, Hilden, Germany) Clean-up kit. The Ribo-Zero treated RNA was quantified by using Agilent Bioanalyzer RNA 6000 Pico Kit (Agilent Technologies, CA, USA) and further used for the cDNA synthesis by using ScriptSeq™ v2 RNA-Seq kit (Epicentre, Illumina, Madison, WI USA). The cDNA was purified using AMPure XP (Beckman Coulter, Brea, CA, USA) beads and multiplexed by using ScriptSeq Index PCR Primers (Epicentre, Illumina, Madison, WI USA). cDNA libraries were quantified by using KAPA Illumina Library Quantification kit (KAPA Biosystems, Wilmington, MA). Finally, six libraries, which contains the biological triplicate of S*. maltophilia* ATCC 13637^T^ cultured at 28 °C (SM_28_R1, SM_28_R2, SM_28_R3) and 37 ◻ (SM_37_R1, SM_37_R2, SM_37_R3) were pooled and sequenced using in-house Illumina MiSeq (Illumina, Inc., San Diego, CA, USA) platform with 2 ×75 bp paired end run.

### RNA-Seq data analysis

The indexing adapters were trimmed by MiSeq control software during the base calling and read quality assessment was done using FastQC version 0.11.2 (Andrews, 2010; Babraham Bioinformatics, Cambridge, UK). The complete genome sequences of *S. maltophilia* ATCC 13637^T^ (Accession No: NZ_CP008838) was downloaded from NCBI-GenBank (https://www.ncbi.nlm.nih.gov/genome/880?genome_assembly_id=2052953) and used as a reference for aligning the reads by using Bowtie 2 (Langmead and Salzberg 2012). The aligned SAM files generated by bowtie were sorted using samtools v1.4.1 (Li, Handsaker et al. 2009). The obtained BAM files were used as input to *cufflinks* v2.2.1 (Trapnell, Williams et al. 2010, Trapnell, Roberts et al. 2012, Trapnell, Hendrickson et al. 2013), which was used to assemble transcripts with FPKM (*f*ragments *p*er *k*ilobase of transcript per *m*illion mapped reads) values. The data files for the replicates were merged into single transcript with *Cuffmerge* and differential gene expression analysis between both condition, i.e. 28 ◻ and 37 ◻ was performed using the *Cuffdiff*, a package of the cufflinks v2.2.1 (Trapnell, Williams et al. 2010, Trapnell, Roberts et al. 2012, Trapnell, Hendrickson et al. 2013). The output data from *Cuffdiff* were imported to cummeRbund v2.18.0 (Goff, Trapnell et al. 2013), which is based on R statistical package version 3.4.0 for visualization. Supplementary Figure 2 shows the workflow employed for differential gene expression analysis using RNA-Seq. Gene expression data were deposited to the Gene Expression Omnibus database (accession number: GES101926).

**Figure 2:**
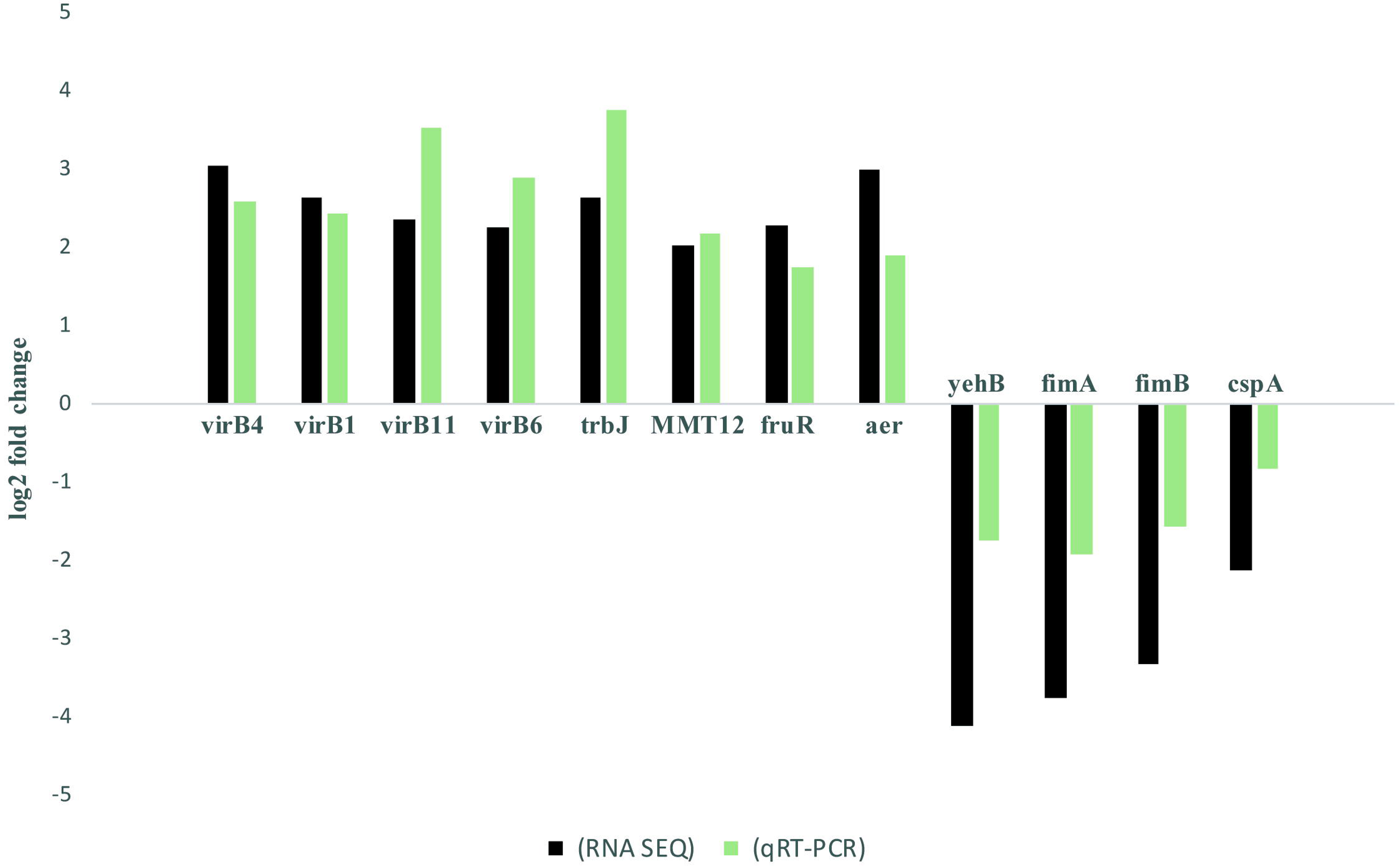
qRT-PCR validation of differentially expressed genes. Expression profile of twelve genes by RNA-Seq and qRT-PCR.

### qRT-PCR validation of the differentially expressed genes

To confirm some of the differential expressed genes obtained using RNA-Seq, a conventional real-time quantitative reverse transcription-PCR (qRT-PCR) was employed to measure changes in the mRNA level of each gene. Gene-specific primers of the differentially expressed genes were designed by using primer3 tool (http://bioinfo.ut.ee/primer3-0.4.0/) and listed in supplementary table 2. RNA was isolated from bacterial cells grown at 28 °C and 37 ◻ as described earlier. The quantitative real-time PCR assay was performed with SuperScript III Platinum SYBR Green One-Step qRT-PCR kit (Thermo Scientific, Wilmington, DE, USA). For each sample, three technical replicates were included, and reactions were set up according to manufacturer’s guidelines. The amplification conditions were: cDNA synthesis 50 ◻ for 45 minutes, initial denaturation at 95 ◻ for 5 minutes, 40 cycles of denaturation at 95 ◻ for 15 seconds followed by annealing at 60 ◻ for 30 seconds and extension at 40 ◻ for 30 seconds. Melting curve analysis confirmed that all PCRs amplified a single product. Gene expression levels were normalized to 16S rRNA gene and *ftsZ* gene. The relative expression of each gene at 37 ◻ relative to 28 ◻ was expressed as fold change calculated by using 2^-ΔΔct^ method. The resulting fold change values were converted to log_2_ fold value and were plotted against the log_2_fold of RNA-Seq data.

### Functional categorization of differentially expressed genes

eggNOG v4.5.1, a database (Huerta-Cepas, Szklarczyk et al. 2015) of orthologous groups and functional annotation was used to classify genes differentially expressed at 28 ◻ and 37 ◻ into functional categories based on Clusters of Orthologous Groups (COG).

All the full length differentially expressed genes obtained from the RNA-Seq experiment were fetched from all the type strains of genus *Stenotrophomonas maltophilia* complex (Smc complex) using tBlastn (Camacho, Coulouris et al. 2009). Cut-off for similarity was set to be 60% and coverage was 50%. All the differentially expressed genes from reference genome ATCC 13637^T^ were annotated using eggNOG-mapper v2 (Huerta-Cepas, Szklarczyk et al. 2015). Based on the presence and absence of the gene a heatmap was constructed using GENE-Ev3.0.215 (https://www.broadinstitute.org/).

### Transmission electron microscopy

Transmission electron microscopy was used to visualize the morphology of the flagella at 28 ◻ and 37 ◻. Bacterial cultures were grown in 20ml LB and incubated at 28 ◻ and 37 ◻ respectively until OD600nm reaches to 0.8. The cells were harvested by centrifugation at 2000 rpm for 10 minutes. The cell pellet was washed twice with 1X PBS (Invitrogen, Carlsbad, CA, USA) and finally suspended in 50μl of 1X PBS (Invitrogen, Carlsbad, CA, USA). 10-20μl of bacterial suspension was placed on a carbon-coated copper grid (300 mesh, Nisshin EM Co., Ltd.) for 15 minutes. The grid was then negatively stained for 30 seconds with 2% phosphotungstic acid, dried and examined using JEM 2100 transmission electron microscope (JEOL, Tokyo, Japan) operating at 200 kV.

### Motility Assays

Motility patterns of *Stenotrophomonas maltophilia* ATCC 13637^T^ were assessed by using motility media. For swimming motility, 5μl of overnight grown culture was spotted on plates containing 1% tryptone, 0.5% NaCl and 0.3% agar. Similarly, for swarming motility 5μl of overnight grown culture was spotted on plates containing 1% tryptone, 0.5% NaCl and 0.5% agar. Plates were incubated at 28 ◻ and 37 ◻ for 7 days. Twitching motility was evaluated on plates containing 1% tryptone, 0.5% NaCl and 1.2% agar. A bacterial colony was stabbed deep into the agar to the bottom with the help of a sterile toothpick. Plates were incubated at 28 ◻ and 37 ◻ for 7 days. Then, to check twitching motility, agar was removed, and plates were stained with 0.1 % crystal violet. Motility assays were carried out on three biological replicates.

### Growth curve measurements

The growth curves at two temperatures i.e. 28 ◻ and 37 ◻ was generated by growing bacterial culture at 28 ◻ and 37 ◻ overnight. 1% of the overnight grown culture (OD=0.8 − 1.0) was then inoculated in fresh 50ml LB with an initial OD600nm 0.015. Readings were taken every 1 hour for 32 hours at OD_600_nm.

## Results and Discussion

### Comparative transcriptome analyses of Smal during growth at 28 ◻ and 37 ◻

To determine the genetic mechanism underlying adaptation of Smal at human body temperature, we performed RNA-Seq analysis on three biological replicates of Smal grown at 28 ◻ and 37 ◻ (Supplementary Figure 1).

A total 4,676,670, 9,477,113, 7,989,000 and 3,536,078, 11,310,235 and 14241935 sequencing reads were obtained for three biological replicates for growth at 28 ◻ (SM_28_R1, SM_28_R2, SM_28_R3) and 37 ◻ (SM_37_R1, SM_37_R2, SM_37_R3) respectively. Reads from all replicates were mapped to the reference genome *S. maltophilia* ATCC 13637^T^ with overall mapping frequency ranging from 87% to 94% (Supplementary Table 1).

To identify differentially expressed genes at 37 ◻, we compared transcript profiles of *S. maltophilia* ATCC 13637^T^ grown at 28 ◻ and 37 ◻. The global transcriptional profiles for two conditions were obtained by data normalization and statistical analysis (Supplementary Figure 2). A matrix of pairwise comparison based on the FPKM (Fragments Per Kilobase of transcript per Million mapped reads) values between two conditions was obtained. It was used to generate the volcano plot (Figure 1A) to map the fold change in transcript expression against its statistical significance (p-values).

Total 51 genes were differentially expressed when the *S. maltophilia* ATCC 13637^T^ was grown at 37 ◻ as compared to growth at 28 ◻ with the statistically significant cut off values: *p*-value < 0.05, *q*-value < 0.001 and log_2_ fold change > 2. Among differentially expressed genes, 13 genes (accounting for the 25% of differentially expressed genes) were upregulated (Table 1) while 38 genes (accounting for 75%) were downregulated at 37 ◻ as compared to the 28 ◻ (Table 2). The classification of differentially expressed genes by COG (cluster of orthologous groups) revealed that genes in sixteen COG classes were differentially expressed (Figure 1 B). The most COG categories for which the greater number of the genes were differentially expressed are intracellular trafficking and secretion, signal transduction, cell motility and with unknown function (Figure 1 B). The differentially expressed genes belonging to the cell motility; secondary structure; post-translational modification; replication and repair; translation; lipid metabolism; coenzyme metabolism; nucleotide metabolism and transport; amino acid metabolism and transport classes were downregulated at 37 ◻.

To partially validate the differentially expressed genes during growth at 37 ◻ as compared to at 28 ◻, we performed the qRT-PCR analysis of selected genes. We have analyzed the expression profiles of randomly selected twelve differentially expressed genes at 37 ◻ and 28 ◻ (Figure 2). We have used 16S rRNA and *ftsZ* genes as internal control. High correlation (R2 = 0.9135) between expression levels of genes measured by the RNA-Seq and qRT-PCR was observed (Figure 2).

### Temperature dependent regulation of cell motility

Majority of the differentially expressed genes belong to the cell motility category, and all of them were downregulated at 37 ◻. These include isoforms of a gene (DP16_RS19060, DP16_RS19065), which encodes for fimbrial outer membrane protein and type I fimbrial proteins, fimbrial proteins (DP16_RS19075), fimbrial chaperone (DP16_RS19070), methyl-accepting chemotaxis (DP16_RS03855), flagellin (DP16_RS11160), methyl-accepting chemotaxis protein (DP16_RS21325), CheV chemotaxis protein (DP16_RS11100) and methyl-accepting chemotaxis DP16_RS22960. The methyl-accepting chemotaxis proteins and CheV chemotaxis protein are categorized into signal transduction class along with GTP-binding protein TypA (DP16_RS17970), histidine kinase (DP16_RS08460), and signal transduction protein with HDOD domain (DP16_RS21040), which were also down regulated (Table 2).

In order to check the phenotypic effect of downregulation of the cell motility and chemotaxis genes at 37 ◻, we have performed the swimming and swarming motility assay during growth at 28 ◻ and 37 ◻. The swimming and swarming motility is affected at 37 ◻ as compared to that of 28 ◻ (Figure 3 A and B). Further, downregulation of genes involved in flagellin biosynthesis leads to the development of less or impaired flagella at 37 ◻ as compared to the 28 ◻, which was observed in transmission electron microscopy. (Figure 3 C). The impaired flagella ultimately affect the motility at 37 ◻ as compared to the 28 ◻. Taken together, these observations suggest the thermoregulation of cell motility in *S. maltophilia*.

**Figure 3:**
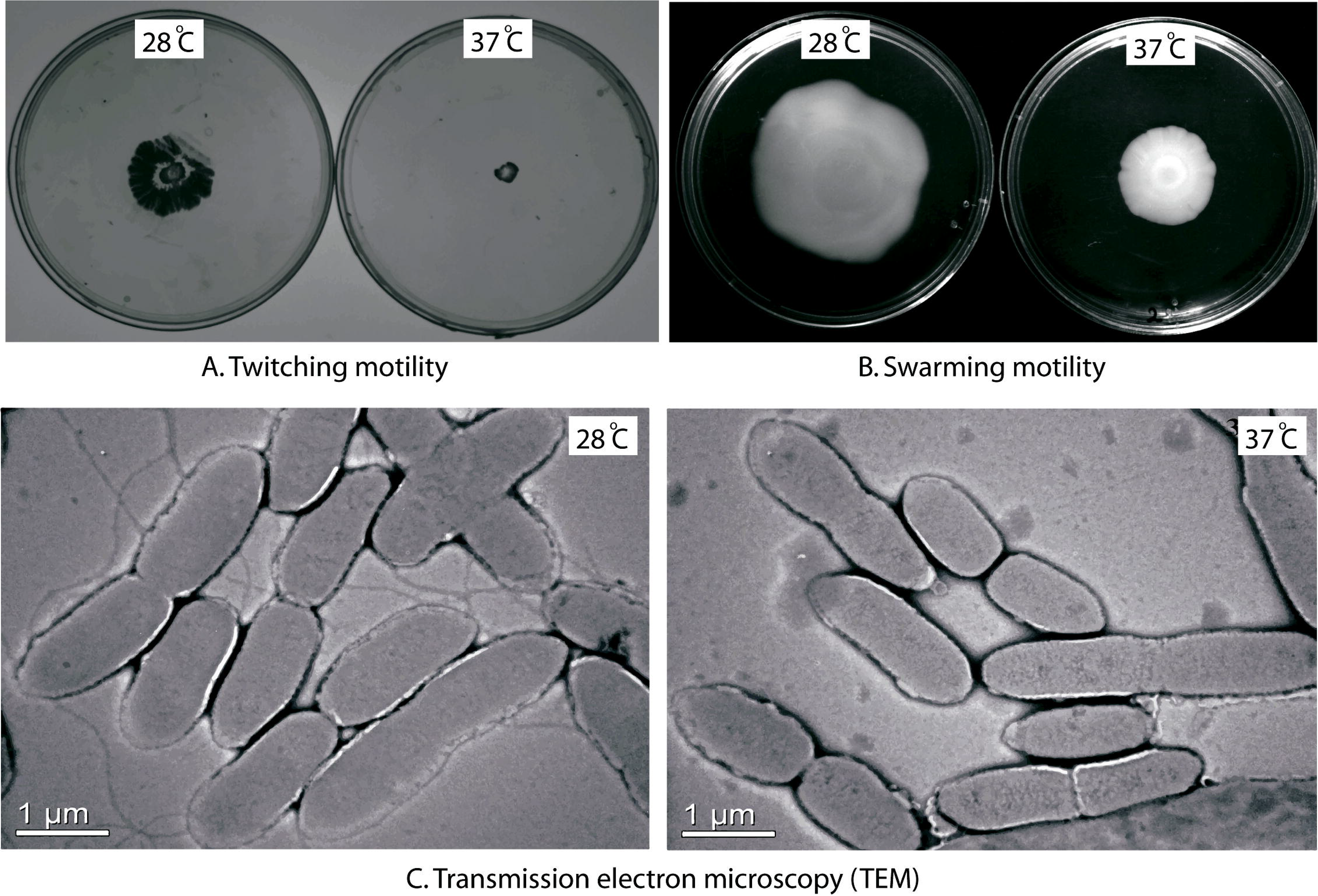
Temperature-dependent regulation of motility. A) Twicthing motility of *S. maltophilia* ATCC 13637^T^ observed during growth at 28 ◻ and 37 ◻. B) Swarming motility of *S. maltophilia* ATCC 13637^T^ observed at growth 28◻ and 37◻. C) Transmission electron micrographs of *S. maltophilia* ATCC 13637^T^ grown at 28 ◻ and 37 ◻ on nutrient agar and negatively stained with 1% phosphotungstic acid.

In bacterial pathogens, it is now a well-known fact that virulence-related traits are generally overexpressed at physiological temperature, i.e. 37 ◻ (Konkel and Tilly 2000). The repression of motility genes at 37 °C to avoid the host recognition was also reported in *Listeria monocytogenes*, which is a foodborne pathogen of environmental origin (Gründling, Burrack et al. 2004). In *Listeria monocytogenes, mogR* transcriptional repressor of flagellar genes along with a protein thermometer *gmaR* which represses flagellar biosynthesis at 37 ◻. The temperature dependent regulation of motility was observed in several human and plant pathogens like *Yersinia enterocolitica, Listeria monocytogenes, Escherichia coli* and *Pseudomonas syringae* (Kapatral, Olson et al. 1996, Kamp and Higgins 2011, Hockett, Burch et al. 2013, Sciandrone, Forti et al. 2019).

The flagella and fimbriae serve as pattern recognition molecule (PAMP), which activate innate immune response in the host cell, thus act as an essential virulence factor for Smal (Zgair and Chhibber 2010). Despite the important role of flagellin and fimbria genes in the Smal pathogenesis, these genes were downregulated at 37 ◻ suggesting that it is an adaptive mechanism by which Smal avoids host recognition and subsequent host innate immune response. In Smal *FsnR* is a canonical positive regulator directly or indirectly controlling the transcription of most flagellar genes by binding to the promoter region of the flagellar biosynthesis gene cluster (Kang, Wang et al. 2015). There might be an involvement of the unidentified protein thermometer, which along with the FsnR may regulate the temperature-dependent flagellar motility. The chemotaxis involves selective movements by using flagella and pili towards nutrients or to escape from hostile environments. There is downregulation of the multiple key genes involved in chemotaxis, which is in accordance with the downregulation of the flagellin genes.

### Downregulation of genes involved in energy production, metabolism and protein synthesis

The expression of two genes involved in energy production and conservation were downregulated at 37 ◻. These include the ATP synthase subunit beta (DP16_RS01805) and C4-dicarboxylate transporter (DP16_RS01020) responsible for uptake of fumarate, succinate and malate, which are essential intermediates in TCA cycle. Apart from this, there is also downregulation of genes belonging to translation, amino acid metabolism and transport, replication and repair, inorganic ion and transport metabolism lipid metabolism, coenzyme metabolism was observed at 37 ◻. Down-regulated genes belong to inorganic ion transport and metabolism category, including phosphate-selective porin O and P (DP16_RS01055), iron uptake factor (DP16_RS02290) and protein of unknown function with domain DUF47 (DP16_RS13710). The data suggested that downregulation of two genes involved in translation (DP16_RS15085, DP16_RS00510), which encodes for a protein that removes the N-terminal methionine from nascent proteins and 50S ribosomal protein L31 type b. The genes belonging to COG class: Post-translational modification, protein turnover, chaperone functions DP16_RS20730 (peptidylprolyl cis-trans isomerase), nucleotide metabolism and transport, DP16_RS12245 (ribosome biogenesis GTPase), amino acid metabolism and transport, DP16_RS02690 (S-adenosylmethionine decarboxylase proenzyme), DP16_RS02840 (Dihydroorotate dehydrogenase), Energy production and conversion DP16_RS01020 (Sodium dicarboxylate symporter family), DP16_RS01805 (ATP synthase subunit B) were also downregulated. The isoforms of the genes DP16_RS20765 (Exodeoxyribonuclease 7 small subunit) and DP16_RS20770 (Polyprenyl synthetase) belonged to COG classes replication and repair, coenzyme metabolites respectively were downregulated at 37 ◻. The downregulation of genes involved in energy production and metabolism; translation is reflected in the lower growth rate of *S. maltophilia* ATCC 13637^T^ at 37 ◻ as compared to 28 ◻ (Supplementary Figure 3). This also suggests a reduction in energy production processes in *S. maltophilia* ATCC 13637^T^ may represent a survival strategy during adaptation at human body temperature.

### Upregulation of VirB/D4 Type IV secretion system at 37 ◻

Comprehensive functional and COG analyses of upregulated genes revealed that five pivotal genes DP16_RS07185/ *virB4* (log2 FC=3.04), DP16_RS07180/*trbJ* (log2 FC=2.6), DP16_RS07200/ *virB1* 3 (log2 FC=2.6), DP16_RS07205/ *virB11* (log2 FC=2.34) and DP16_RS07175/*virB6* (log2 FC=2.2) that are part of Type IV VirB/D4 secretory system, were upregulated at 37 ◻ (Table 1). The expression of the VirB/D4 T4SS components *virB4*, *trbJ, virB1, virB11* and *virB6* was higher at 37 ◻ suggesting that VirB/D4 T4SS in *S. maltophilia* ATCC 13637^T^ is regulated by the temperature. T4SS in Smal is horizontally acquired and present on the genomic island. It is present in the eight other non-clinical species of genus *Stenotrophomonas*, *i.e., S. chelatiphaga, S. daejeonensis, S. ginsengisoli, S. indicatrix, S. koreensis, S. lactitubi, S. pavanii, and S. pictorum* (Nas, White et al. 2019). The VirB/D4 T4SS is absent in the *S. acidaminiphila, S. nitritireducens, S. panacihumi, S. rhizophila, and S. terrae* (Nas, White et al. 2019). Apart from the role in conjugation, T4SSs also play an important in the pathogenic mechanism of many animal pathogens *Legionella pneumophila*, *Bordetella pertussis*, *Coxiella burnetii*, *Bartonella henselae*, *Brucella* spp. and *Helicobacter pylori* as well as plant pathogen *Agrobacterium tumefaciens* (Souza, Oka et al. 2015, Gonzalez-Rivera, Bhatty et al. 2016). VirB/D4 T4SS of *S. maltophilia* is related to the well-known plant pathogens of *Xanthomonas* species, a phylogenetic relative of Smal, which mediates killing of the other bacterial cell by T4SS but not involved in virulence (Souza, Oka et al. 2015). In the latest study by Nas et al. suggested that VirB/D4 T4SS in Smal inhibits the apoptosis in an epithelial cell to enhance attachment while it promotes apoptosis in infected mammalian macrophages to escape from phagocytosis (Nas, White et al. 2019). The study further revealed that VirB/D4 T4SS in Smal stimulates the growth and mediates inter bacterial killing of other bacteria in the complex microbial community (Nas, White et al. 2019). Thus, by considering the role of VirB/D4 T4SS in virulence, adaptation in the complex microbial community and its upregulation at 37 ◻ suggests a temperature-dependent strategy for pathoadaption.

### Upregulation of the genes involved in the aerotaxis, cation diffusion facilitator family transporter and LacI family transcriptional regulators

Interestingly, increased expression of genes involved in aerotaxis, which is also known as energy taxis at 37 ◻. It is a behavioural response that guides bacterial cells to navigate toward micro-environments where oxygen concentration, energy sources, and redox potential are optimal for growth (Taylor, Zhulin et al. 1999). This process is coordinated by aerotaxis receptor *Aer*, which measures redox potential. It infers energy levels *via* a flavin adenine dinucleotide (FAD) cofactor bound to a cytoplasmic PAS domain (Taylor and Zhulin 1999, Campbell, Watts et al. 2011). In *S. maltophilia* ATCC 13637^T^, two genes (DP16_RS19060 and DP16_RS19065) that encode for FAD-binding domain protein and PAS sensor domain-containing protein (Bouckaert, Heled et al.) are transcribed as single transcript and are overexpressed at 37 ◻. This may help Smal to adapt and colonize different niches with a different oxygen gradient. Thus, further experiments are required to understand the role of aerotaxis in Smal adaptation to human host and virulence. Reports are citing the role of aerotaxis in an adaptation of *C. jejuni* at human gut with different oxygen gradient and in *Ralstonia solanacearum* it is required for the biofilm formation (Hazeleger, Wouters et al. 1998, Yao and Allen 2007). The role of the aerotaxis in virulence of bacteria is not fully understood, but it plays an important role in the adaptation of bacterium toward its host (Henry and Crosson 2011).

Among the upregulated gene, DP_RS06915 (log2 FC=2.0), that code for cation diffusion facilitator (CDF) family transporter is important for the transition of metals efflux from the cytosol to periplasm. CDF transporter plays a role in the transition metal tolerance, i.e., exporting metal surplus from cell to avoid excessive accumulation and toxicity. Apart from the role in the transition metals efflux, they also participate in the infection process in *P. aeruginosa* (Salusso and Raimunda 2017). As the transcription of CDF was increased at 37 ◻ and by considering its possible role in the infection process, it is necessary to assess the role of CDF in virulence and adaption of Smal.

The transcription regulator of LacI family DP16_RS10420 is overexpressed (log2 FC=2.2) at 37 ◻. This family of transcriptional regulators is known to play an essential role in the carbohydrate uptake or metabolism and virulence (Van Gijsegem, Wlodarczyk et al. 2008, Njoroge, Nguyen et al. 2012, Ravcheev, Khoroshkin et al. 2014). Upregulation of the gene *fruR*, which is a transcription factor and belongs to the LacI family was observed at 37 °C, suggesting it may play an important role in adaptation and virulence. Two genes containing the domain of unknown function DUF 4189 (DP16_RS23790, DP16_RS23785) and one gene that encode a hypothetical protein were upregulated at 37 ◻. Isoforms of the genes DP16_RS19060 and DP16_RS19065 which codes for sulphite reductase subunit alpha with FAD-binding domain and PAS sensor domain-containing protein is also overexpressed (log2 FC=2.9) at 37 °C. Therefore, future studies are needed to reveal the role of these genes in infection and adaptation to human body temperature.

### Human body temperature is not heat stress for Smal

From differential expression analysis of Smal at two temperatures, we observed a significant downregulation of gene for cold shock protein (*cspA2*), which belongs to transcription COG class is downregulated (log2 FC= −2.1) at 37 ◻ suggesting its role in adaptation to lower environmental temperature. Despite the presence of heat shock chaperons in Smal, we did not find differential gene expression of heat shock response genes, which is generally indicative of heat stress. This suggests that Smal has evolved to thrive at human body temperature without a need to activate protective surveillance responses against heat stress. Overall, this emphasizes that human body temperature is not heat stress for Smal. This kind of response was also reported in the environmentally originated opportunistic pathogen *Pseudomonas aeruginosa* during growth at 37 ◻ (Wurtzel, Yoder-Himes et al. 2012).

In addition to clinical, *S. maltophilia* complexes have species from diverse lifestyles. Here, we looked for status of all the 49 pathoadaptive or differentially expressed genes status in all the species of Smc (Figure 4). Interestingly, ShlB/FhaC/HecB family hemolysin secretion/activation protein (DP16_RS23075) and hypothetical protein (DP16_RS23790) are unique to Sma which can be correlated with its clinical lifestyle. While, motility and T4SS genes in few of the other species genus *Stenotrophomonas.* Energy production, metabolism and transcription regulators are largely present in all the species of Smc. Overall phylogenomic based transcritomic understanding reveals that the transistion and success of S. maltophilia species in the genus has been intricate by modulating functions related to immune evasion as seen by downregulation of flagella, protection from host defense responses as seen by downregulation of genes involved in motility apart from other cellular processes related to physiology, replication and transcription (Figure 5). Further molecular genetic studies on the differentially expressed that are unique to Sma may allow understanding success of this species as opportunistic human pathogen.

**Figure 4:**
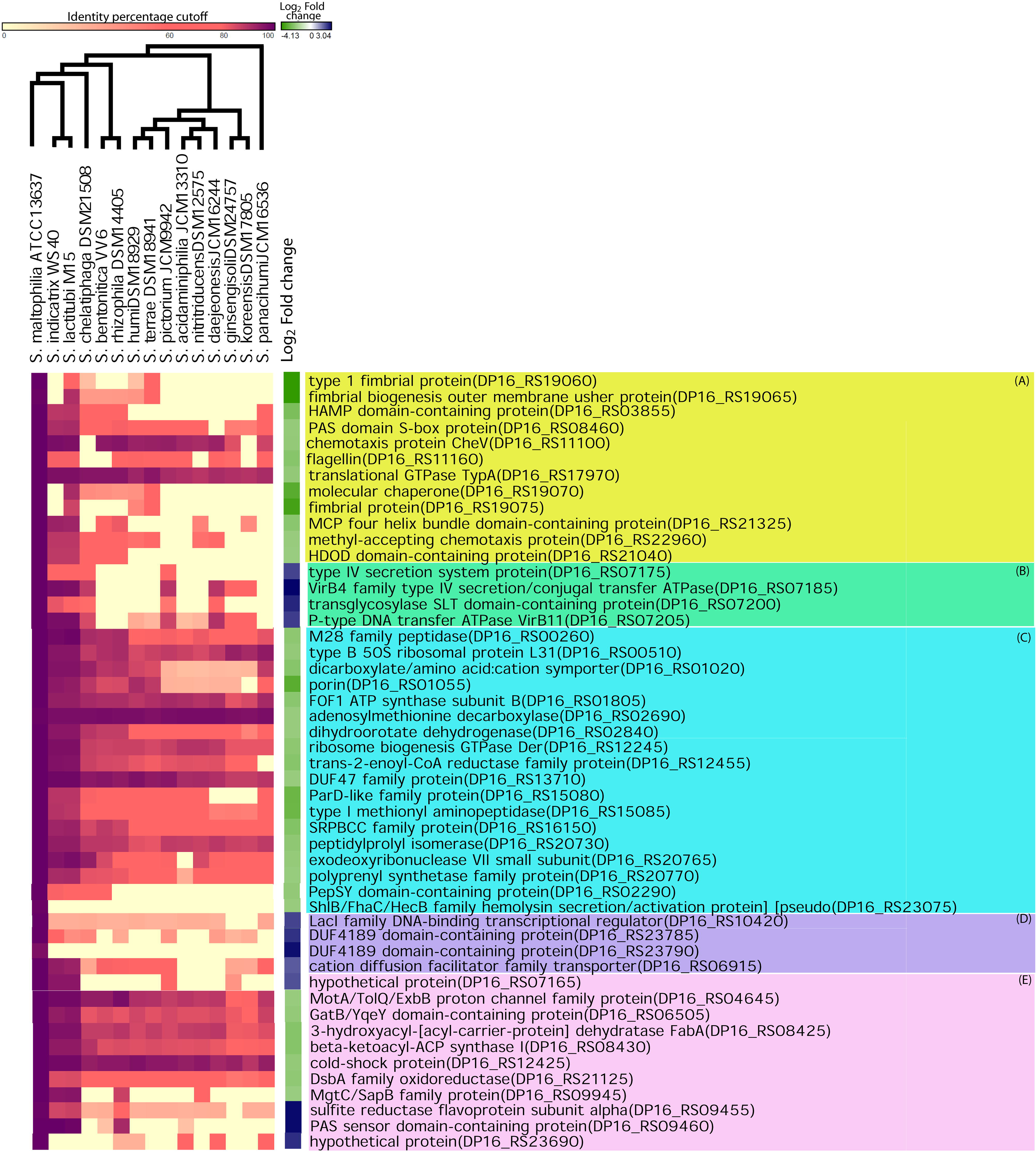
Heatmap showing presence or absence of differentially expressed genes in Smc along with log_2_ fold change of the genes at 37 ◻ as compared to 28 ◻. Genes related to (A) motility, (B) type IV secretion system, (C) energy, metabolism (D) aerotaxis, cation diffusion facilitator family transporter and LacI family transcription regulators and (E) others.

**Figure 5:**
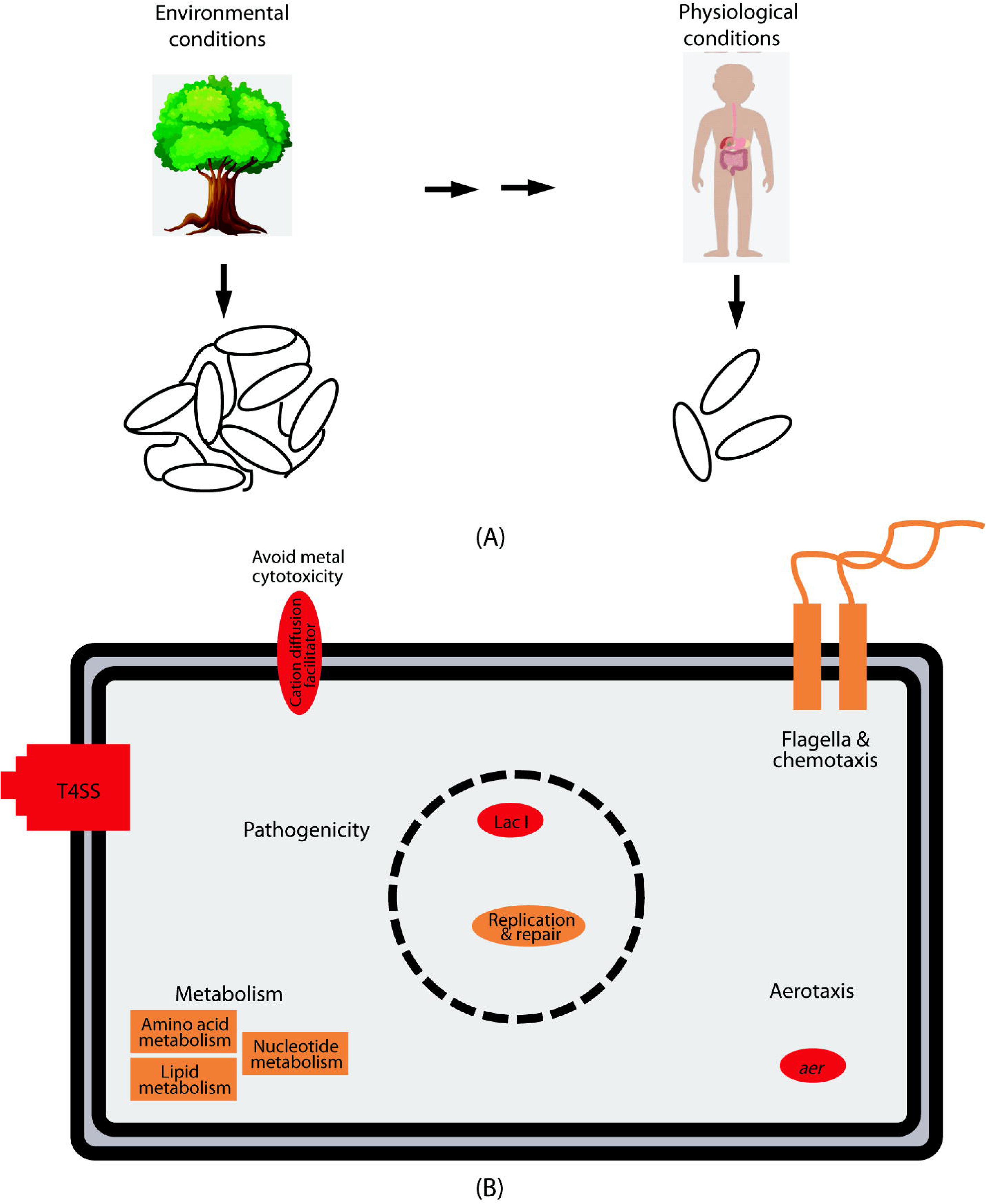
**(A)**Transition of Smal from environment to clinical settings **(B)**Schematic diagram of upregulated (purple) and downregulated (green) genes.

## Conclusion

Current work is a high-resolution comprehensive comparative analysis of RNA-Seq based transcriptome of opportunistic pathogen *S. maltophilia.* This study has provided a framework for studying the molecular mechanism underlying transition of an environmental bacterium to become a successful human pathogen. The study also suggests how *S. maltophilia* is a matter of grave concern to the immunocompromised patient. Further, studies on the characterization of differentially expressed genes of *S. maltophilia* at physiological temperature will give more insights into its adaptation to human host and pathogenesis.

## Author statements

### Authors and contributors

PPP, SM, SK has prepared the RNA-Seq library preparation and performed transcriptome sequencing on Illumina MiSeq platform. PPP, SK and KB have performed computational analysis of RNA-Seq data analysis. PPP and AK has performed motility assay, growth curve and validation of differentially expressed genes by RT-qPCR. AK, SK and PPP performed transmission electron microscopy (TEM). PPP has drafted the manuscript with inputs from SK, KB and AK. PBP and PPP have conceived and designed the experiments with inputs from SK, AK and KB.

## Conflicts of interest

The authors declare that there are no conflicts of interest.

## Funding information

This work is supported by Research Council approved projects of CSIR-IMTECH, OLP-148 and OLP-601 to PBP. All authors declare no conflict of interests. PPP is acknowledging Senior Research Fellowship from University Grant Commission, New Delhi. SM and SK fellowship from Council of Scientific and Industrial Research, India. AK is supported by Senior Research Fellowship from INSPIRE scheme of the Department of Sciences and technology, Government of India.

---

**Table 1:** *S. maltophilia* ATCC 13637^T^ genes significantly up-regulated during the growth at 37 ◻.

**Table 2:** *S. maltophilia* ATCC 13637^T^ genes significantly down-regulated during growth at 37 ◻.

## Figures and Tables Legend of Supplementary files

**Supplementary Figure 1: Experimental workflow for differential gene expression analysis of *S. maltophilia* grown at 28 ◻ and 37**◻. R1, R2 and R3 denote the biological replicates.

**Supplementary Figure 2: Workflow employed for RNA-Seq data analysis for differential gene expression:**First quality reads from both conditions were mapped to the reference genome with Bowtie2. The mapping was done independently for each read from biological replicates. The mapped SAM files were converted to sorted BAM files using SAMtools. Sorted BAM files were given as input to *Cufflinks*, which produces unary assembled transcripts for each replicate. The assembly files were merged with reference transcriptome annotation into a unified annotation using *Cuffmerge* and used for further analysis. This merged annotation was quantified in each condition by *Cuffdiff*, which produces expression data in a set of tabular files. These files were

**Supplementary Figure 3: Growth curve measurements:** Growth curves of *S. maltophilia* ATCC 137637 at two temperatures, i.e., 28 ◻ and 37 ◻

**Supplementary Table 1: Summary of Illumina RNA-Seq data generated.***S. maltophilia* ATCC 13637 grown at 28 ◻ (SM_28) and 37 ◻ (SM_37) number (_1, _2, _3) following SM_28 and SM_37 represents replicates for each condition.

**Supplementary Table 2: List of primers used in qRT-PCR for validation of RNA-Seq.**

## Notes

### Competing Interest Statement

The authors have declared no competing interest.

## References

Alavi, P., et al. (2014). “Stenotrophomonas comparative genomics reveals genes and functions that differentiate beneficial and pathogenic bacteria.” BMC genomics 15(1): 482.

Bansal, K., et al. (2020). “Deep phylo-taxono-genomics (DEEPT genomics) reveals misclassification of Xanthomonas species complexes into Xylella, Stenotrophomonas and Pseudoxanthomonas.” BioRxiv.

Bouckaert, R., et al. (2014). “BEAST 2: a software platform for Bayesian evolutionary analysis.” PLoS computational biology 10(4): e1003537.

Brooke, J. S. (2012). “Stenotrophomonas maltophilia: an emerging global opportunistic pathogen.” Clinical microbiology reviews 25(1): 2–41.

Brooke, J. S. (2014). New strategies against Stenotrophomonas maltophilia: a serious worldwide intrinsically drug-resistant opportunistic pathogen, Taylor & Francis.

Camacho, C., et al. (2009). “BLAST+: architecture and applications.” BMC bioinformatics 10(1): 421.

Campbell, A. J., et al. (2011). “Role of the F1 region in the Escherichia coli aerotaxis receptor Aer.” Journal of bacteriology 193(2): 358–366.

Creecy, J. P. and T. Conway (2015). “Quantitative bacterial transcriptomics with RNA-seq.” Current opinion in microbiology 23: 133–140.

Croucher, N. J. and N. R. Thomson (2010). “Studying bacterial transcriptomes using RNA-seq.” Current opinion in microbiology 13(5): 619–624.

Goff, L., et al. (2013). “cummeRbund: Analysis, exploration, manipulation, and visualization of Cufflinks high-throughput sequencing data.” R package version 2(0).

Gonzalez-Rivera, C., et al. (2016). “Mechanism and function of type IV secretion during infection of the human host.” Microbiology spectrum 4(3).

Gründling, A., et al. (2004). “Listeria monocytogenes regulates flagellar motility gene expression through MogR, a transcriptional repressor required for virulence.” Proceedings of the National Academy of Sciences of the United States of America 101(33): 12318–12323.

Hazeleger, W. C., et al. (1998). “Physiological activity of Campylobacter jejuni far below the minimal growth temperature.” Applied and Environmental Microbiology 64(10): 3917–3922.

Henry, J. T. and S. Crosson (2011). “Ligand-binding PAS domains in a genomic, cellular, and structural context.” Annual review of microbiology 65: 261–286.

Hockett, K. L., et al. (2013). “Thermo-regulation of genes mediating motility and plant interactions in Pseudomonas syringae.” 8(3): e59850.

Huerta-Cepas, J., et al. (2015). “eggNOG 4.5: a hierarchical orthology framework with improved functional annotations for eukaryotic, prokaryotic and viral sequences.” Nucleic acids research 44(D1): D286–D293.

Kamp, H. D. and D. E. J. P. P. Higgins (2011). “A protein thermometer controls temperature-dependent transcription of flagellar motility genes in Listeria monocytogenes.” 7(8): e1002153.

Kang, X.-M., et al. (2015). “Genome-wide identification of genes necessary for biofilm formation by nosocomial pathogen Stenotrophomonas maltophilia reveals that orphan response regulator FsnR is a critical modulator.” Appl. Environ. Microbiol. 81(4): 1200–1209.

Kapatral, V., et al. (1996). “Temperature◻dependent regulation of Yersinia enterocolitica class III flagellar genes.” 19(5): 1061–1071.

Konkel, M. E. and K. Tilly (2000). “Temperature-regulated expression of bacterial virulence genes.” Microbes and Infection 2(2): 157–166.

Kortmann, J. and F. Narberhaus (2012). “Bacterial RNA thermometers: molecular zippers and switches.” Nature reviews. Microbiology 10(4): 255.

Kumar, S., et al. (2019). “Phylogenomics insights into order and families of Lysobacterales.” Access Microbiology 1(2): e000015.

Lam, O., et al. (2014). “Thermal control of virulence factors in bacteria: A hot topic.” Virulence 5(8): 852–862.

Langmead, B. and S. L. Salzberg (2012). “Fast gapped-read alignment with Bowtie 2.” Nature methods 9(4): 357–359.

Li, H., et al. (2009). “The sequence alignment/map format and SAMtools.” Bioinformatics 25(16): 2078–2079.

Looney, W. J., et al. (2009). “Stenotrophomonas maltophilia: an emerging opportunist human pathogen.” The Lancet infectious diseases 9(5): 312–323.

Mukherjee, P. and P. Roy (2016). “Genomic potential of Stenotrophomonas maltophilia in Bioremediation with an Assessment of its Multifaceted Role in Our Environment.” Frontiers in microbiology 7.

Nas, M. Y., et al. (2019). “Stenotrophomonas maltophilia Encodes a VirB/D4 Type IV Secretion System That Modulates Apoptosis in Human Cells and Promotes Competition Against Heterologous Bacteria Including Pseudomonas aeruginosa.” Infection and Immunity: IAI. 00457–00419.

Njoroge, J. W., et al. (2012). “Virulence meets metabolism: Cra and KdpE gene regulation in enterohemorrhagic Escherichia coli.” MBio 3(5): e00280–00212.

Patil, P. P., et al. (2018). “Taxonogenomics reveal multiple novel genomospecies associated with clinical isolates of Stenotrophomonas maltophilia.” Microbial genomics 4(8).

Patil, P. P., et al. (2016). “Genome sequence of type strains of genus Stenotrophomonas.” Frontiers in microbiology 7: 309.

Ravcheev, D. A., et al. (2014). “Comparative genomics and evolution of regulons of the LacI-family transcription factors.” Frontiers in microbiology 5.

Ryan, R. P., et al. (2009). “The versatility and adaptation of bacteria from the genus Stenotrophomonas.” Nature reviews. Microbiology 7(7): 514.

Salusso, A. and D. Raimunda (2017). “Defining the Roles of the Cation Diffusion Facilitators in Fe2+/Zn2+ Homeostasis and Establishment of their Participation in Virulence in Pseudomonas aeruginosa.” Frontiers in cellular and infection microbiology 7.

Sciandrone, B., et al. (2019). “Temperature-dependent regulation of the Escherichia coli lpxT gene.” 1862(8): 786–795.

Shapiro, R. S. and L. E. Cowen (2012). “Thermal control of microbial development and virulence: molecular mechanisms of microbial temperature sensing.” MBio 3(5): e00238–00212.

Souza, D. P., et al. (2015). “Bacterial killing via a type IV secretion system.” Nature communications 6: 6453.

Taylor, B. L. and I. B. Zhulin (1999). “PAS domains: internal sensors of oxygen, redox potential, and light.” Microbiology and Molecular Biology Reviews 63(2): 479–506.

Taylor, B. L., et al. (1999). “Aerotaxis and other energy-sensing behavior in bacteria.” Annual Reviews in Microbiology 53(1): 103–128.

Trapnell, C., et al. (2013). “Differential analysis of gene regulation at transcript resolution with RNA-seq.” Nature biotechnology 31(1).

Trapnell, C., et al. (2012). “Differential gene and transcript expression analysis of RNA-seq experiments with TopHat and Cufflinks.” Nature protocols 7(3): 562.

Trapnell, C., et al. (2010). “Transcript assembly and quantification by RNA-Seq reveals unannotated transcripts and isoform switching during cell differentiation.” Nature biotechnology 28(5): 511–515.

Van Gijsegem, F., et al. (2008). “Analysis of the LacI family regulators of Erwinia chrysanthemi 3937, involvement in the bacterial phytopathogenicity.” Molecular plant-microbe interactions 21(11): 1471–1481.

White-Ziegler, C. A., et al. (2007). “Human body temperature (37 C) increases the expression of iron, carbohydrate, and amino acid utilization genes in Escherichia coli K-12.” Journal of bacteriology 189(15): 5429–5440.

Wurtzel, O., et al. (2012). “The single-nucleotide resolution transcriptome of Pseudomonas aeruginosa grown in body temperature.” PLoS pathogens 8(9): e1002945.

Yao, J. and C. Allen (2007). “The plant pathogen Ralstonia solanacearum needs aerotaxis for normal biofilm formation and interactions with its tomato host.” Journal of bacteriology 189(17): 6415–6424.

Zgair, A. K. and S. Chhibber (2010). “Stenotrophomonas maltophilia flagellin induces a compartmentalized innate immune response in mouse lung.” Journal of medical microbiology 59(8): 913–919.

