## Supplementary material for "Global transcriptome analysis of *Stenotrophomonas maltophilia* in response to growth at human body temperature": Supplimentary_Information

**Current Affiliations**

^$^ Department of Microbiology, School of Medicine, University of Washington, Seattle, WA, USA.

^#^ Imperial Life Sciences (P) Limited, Gurgaon, Haryana, 122001, India

^†^Institute of Infection, Veterinary and Ecological Sciences, University of Liverpool, Liverpool, United Kingdom.

***Address for correspondence**

Prabhu B. Patil

Principal Scientist

Bacterial Genomics and Evolution Laboratory

CSIR- Institute of Microbial Technology, Chandigarh, India.

<http://orcid.org/0000-0003-2720-1059>

**Supplementary Figures**

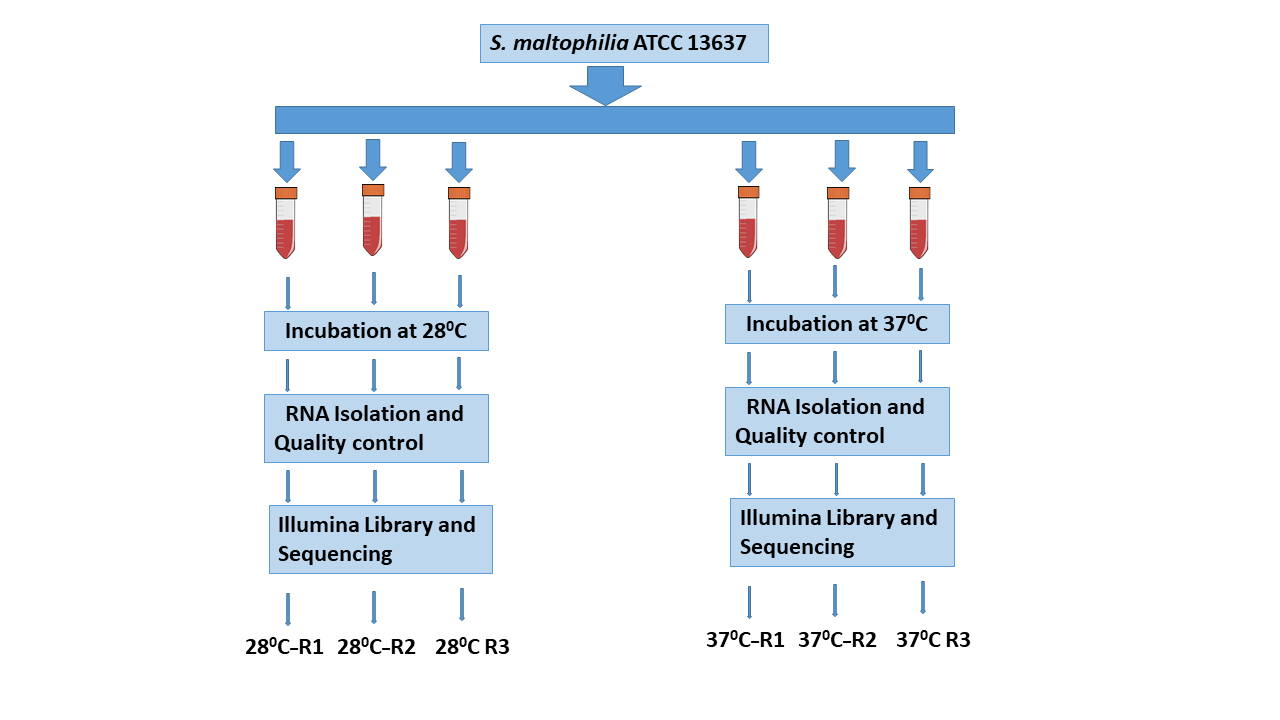

**Supplementary Figure 1: Experimental workflow for differential gene expression analysis of *S. maltophilia* grown at 28 ℃ and 37 ℃.** R1, R2 and R3 denotes the biological replicate.

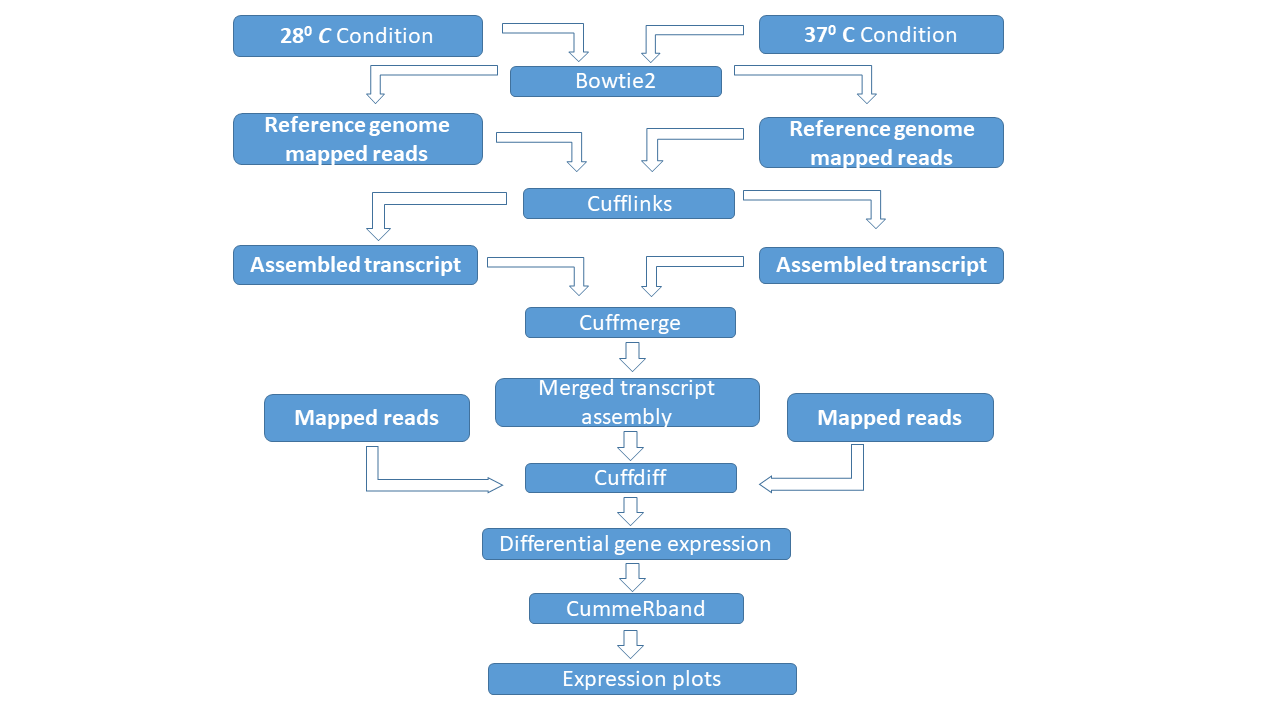

**Supplementary Figure 2: Workflow employed for RNA-Seq data analysis for differential gene expression:** First quality reads from both conditions were mapped to the reference genome with Bowtie2. The mapping was done independently for each reads from biological replicates. The mapped SAM files were converted to sorted BAM files using SAMtools. Sorted BAM files was given as input to *Cufflinks*, which produces unary assembled transcript for each replicate. The assembly files were merged with reference transcriptome annotation into a unified annotation using *Cuffmerge* and used for further analysis. This merged annotation was quantified in each condition by *Cuffdiff*, which produces expression data in a set of tabular files.

**Supplementary Figure 3:**

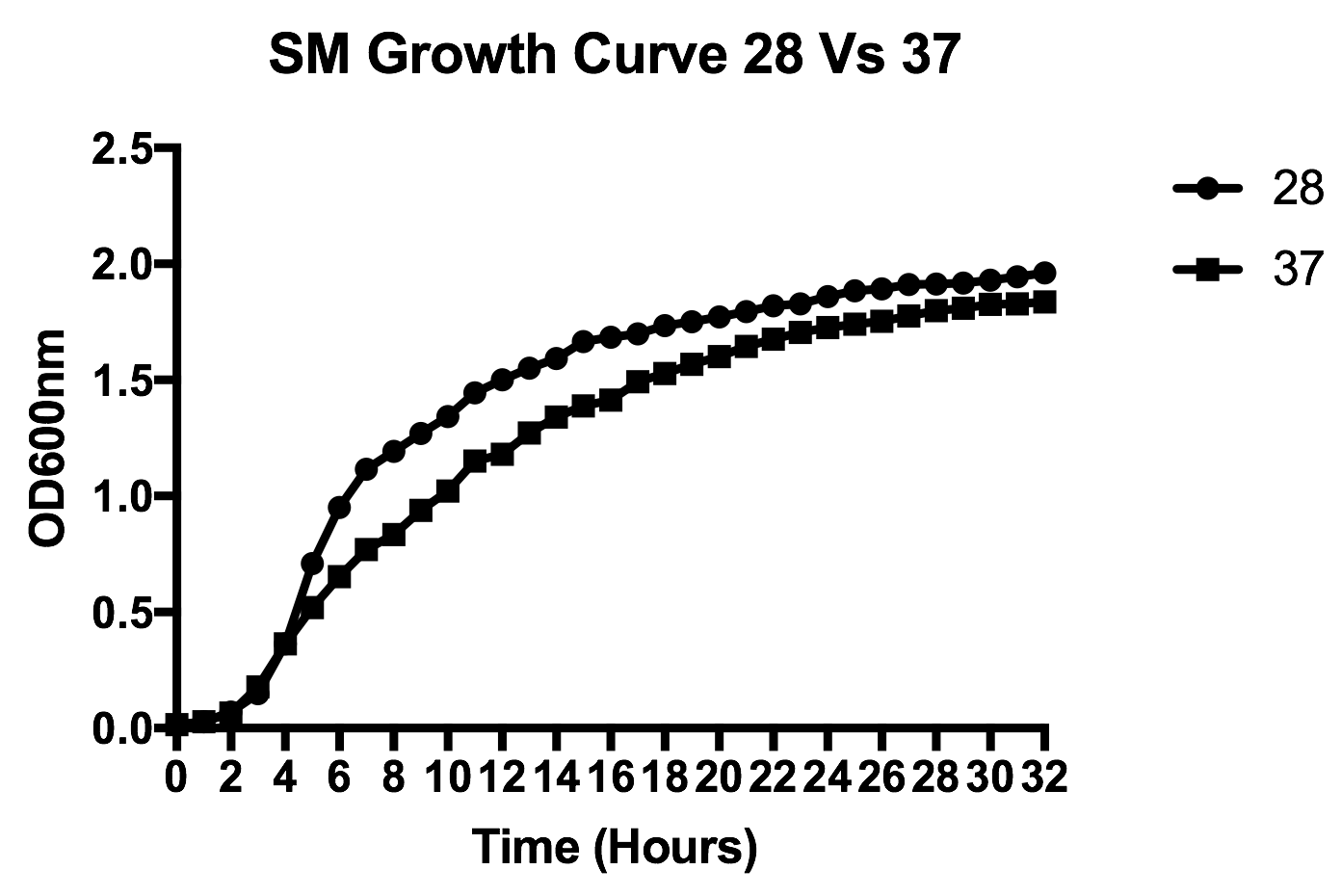

**Supplementary Figure 3: Growth curve measurements:** Growth curves of *S. maltophilia* ATCC 137637 at two temperatures i.e. 28 ℃and 37 ℃.

**Tables:**

**Supplementary Table 1: Summary of Illumina RNA-Seq data generated.** *S. maltophilia* ATCC 13637 grown at 28 ℃ (SM_28) and 37 ℃ (SM_37) number (_1, _2, _3) following SM_28 and SM_37 represents replicates for each condition.

|  | Total quality reads | Total mapped reads | Overall mapping percentage |
| --- | --- | --- | --- |
| SM_28_R1 | 9,353,340 | 8,806,943 | 94.16% |
| SM_28_R2 | 18,954,226 | 17,559,441 | 92.64% |
| SM_28_R3 | 15,978,000 | 14,037,770 | 87.86% |
| SM_37_R1 | 7,072,156 | 6,782,141 | 95.90% |
| SM_37_R2 | 22,620,470 | 22,620,470 | 93.65% |
| SM_37_R3 | 28,483,870 | 28,483,870 | 94.74% |

**Supplementary Table 2:** List of primers used in qRT-PCR for validation of RNA-seq

data

| S.No. | Gene | Primer sequence (5’-3’) |
| --- | --- | --- |
| 1. | *SM-ftsZ-F*  *SM-ftsZ-R* | GGCGCATTTTGAACTGATCG  AGCTTGGCACCGCAATTCT |
| 2. | *SM-fimA-F*  *SM-fimA-R* | TGCCGACCGTGTCCAAGAA  GCACTTGGTCAGGTTGATGG |
| 3. | *SM-fimB-F*  *SM-fimB-R* | ACTCTGGCCGAAGTACATGC  GCCGTAGTCGTTGATGGTGATGAA |
| 4. | *SM-16s-F*  *SM-16s-R* | GACCTTGCGCGATTGAATG  CGGATCGTCGCCTTGGT |
| 5. | *SM-virB1-F*  *SM-virB1-R* | GTCAGGGTCGAACATCATCC  GATGGGTAAACGGTGTAGGC |
| 6. | *SM-virB4-F*  *SM-virB4-R* | TGTGATGGACGAATTCTGGA  ATCACTCTTCAGCGCGTCTT |
| 7. | *SM-virB6-F*  *SM-virB6-R* | GTGCGATGCTGATGCTGTAT  AATGCCGTAGAACAGCCAAC |
| 8. | *SM-virB11-F*  *SM-virB11-R* | CGCGAGTACGCAGAGTTCTT  TCGTTGTCCGGGATATGATT |
| 9. | *SM-trbJ-F*  *SM-trbJ-R* | CATGACATCCCGAAATCACA  GGTCGAAGACAGGGTAACCA |
| 10. | *SM-MMT12-F*  *SM-MMT12-R* | CATCGAAATCCATGTGCTGA  AATCGATGGTCAGCCAGAAC |
| 11. | *SM-yehB-F*  *SM-yehB-R* | CAGTTCAACTCCAGCTTCCTG  ACGTACACGTCGACACGATAGTT |
| 12. | *SM-fruR-F*  *SM-fruR-R* | GATTGTCGAGTACCACGCTGT  CACTGTATCTGCAATTGATGCAC |
| 13. | *SM-aer-F*  *SM-aer-R* | GTATACAAGGACATGTGGGACACC  GATGCTGATGTAGGAGGTGATGT |
| 14. | *SM-cspA2-F*  *SM-cspA2-R* | GGACCTGTTTGTGCACTTCC  GTCAGCCTGCATACCCTTCT |
