## Supplementary material for "Global transcriptome analysis of *Stenotrophomonas maltophilia* in response to growth at human body temperature": Table1

**Table 1:** *S. maltophilia* ATCC 13637^T^ genes significantly upregulated during the growth at 37 ℃ versus 28 ℃.

| Locus tag | Gene description | Gene name | log_2_ Fold change |
| --- | --- | --- | --- |
| DP16_RS07185 | VirB4 family type IV secretion/conjugal transfer ATPase | *virB4* | 3.0401 |
| DP16_RS09455  DP16_RS09460 | PAS sensor domain-containing protein  and sulfite reductase subunit alpha | *-*  *aer* | 2.97384 |
| DP16_RS23790 | DUF4189 domain-containing protein | *-* | 2.92365 |
| DP16_RS07200 | type VI secretion protein | *virB1* | 2.63321 |
| DP16_RS07180 | hypothetical protein | *trbJ* | 2.62841 |
| DP16_RS23690 | hypothetical protein | *-* | 2.5166 |
| DP16_RS23785 | DUF4189 domain-containing protein | *-* | 2.49044 |
| DP16_RS07205 | P-type DNA transfer ATPase VirB11 | *virB11* | 2.34854 |
| DP16_RS10420 | Transcriptional regulator, LacI family | *fruR* | 2.25723 |
| DP16_RS07175 | type VI secretion protein | *virB6* | 2.25345 |
| DP16_RS07165 | hypothetical protein | *-* | 2.12982 |
| DP16_RS06915 | cation transporter | *MMT12* | 2.00271 |
