## Supplementary material for "Global transcriptome analysis of *Stenotrophomonas maltophilia* in response to growth at human body temperature": Table2

**Table 2:** *S. maltophilia* ATCC 13637^T^ genes significantly down regulated during growth at 37 ℃ versus 28 ℃.

| Locus tag | Gene description | Gene name | log_2_ Fold change |
| --- | --- | --- | --- |
| DP16_RS19060  DP16_RS19065 | fimbrial biogenesis outer membrane usher protein/Type 1 fimbrial protein | *mrkD, yehB* | -4.12889 |
| DP16_RS19075 | ferrous iron transporter B | *fimA* | -3.75516 |
| DP16_RS12685 | hypothetical protein | *-* | -3.35666 |
| DP16_RS19070 | fimbrial chaperone | *fimB* | -3.33263 |
| DP16_RS01055 | Porin | *-* | -3.29179 |
| DP16_RS15080 | hypothetical protein | *-* | -2.95503 |
| DP16_RS15085 | type I methionyl aminopeptidase | *map* | -2.93699 |
| DP16_RS03855 | chemotaxis protein | *mcpU* | -2.63073 |
| DP16_RS16150 | polyketide cyclase | *-* | -2.45016 |
| DP16_RS08425,  DP16_RS08430 | beta-hydroxydecanoyl-ACP dehydratase/beta-ketoacyl-[acyl-carrier-protein] synthase I | *fabA, fabB* | -2.40955 |
| DP16_RS11160 | Flagellin | *fliC* | -2.38153 |
| DP16_RS12455 | Short-chain alcohol dehydrogenase family | *-* | -2.36319 |
| DP16_RS01020 | C4-dicarboxylate transporter | *dctA* | -2.35897 |
| DP16_RS21325 | methyl-accepting chemotaxis protein | *mcpU* | -2.30892 |
| DP16_RS01805 | ATP synthase subunit B | *atpF* | -2.30415 |
| DP16_RS12245 | ribosome biogenesis GTPase Der | *der* | -2.27308 |
| DP16_RS00260 | peptidase M28 family protein | *-* | -2.24915 |
| DP16_RS20730 | peptidyl-prolyl cis-trans isomerase | *sylDB* | -2.2321 |
| DP16_RS21125 | DsbA family oxidoreductase | *frnE* | -2.19413 |
| DP16_RS17970 | translational GTPase TypA | *typA* | -2.18648 |
| DP16_RS02290 | Iron-uptake factor | *-* | -2.16518 |
| DP16_RS06505 | transamidase GatB domain protein | *yqeY* | -2.15677 |
| DP16_RS11100 | chemotaxis protein CheV | *cheV2* | -2.14743 |
| DP16_RS00510 | 50S ribosomal protein L31 type B | *rpmE2* | -2.14705 |
| DP16_RS12425 | cold-shock protein | *cspA2* | -2.1415 |
| DP16_RS08460 | hybrid sensor histidine kinase/response regulator | *-* | -2.12618 |
| DP16_RS20765  DP16_RS20770 | Exodeoxyribonuclease 7 small subunit/ (2E,6E)-farnesyl diphosphate synthase | *xseB, ispA* | -2.11889 |
| DP16_RS02690 | S-adenosylmethionine decarboxylase proenzyme | *speD* | -2.08431 |
| DP16_RS22960 | chemotaxis protein | *mcpU* | -2.07516 |
| DP16_RS23075 | Hypothetical protein | *fhaC* | -2.06461 |
| DP16_RS04645 | Biopolymer transporter ExbB | *exbB* | -2.06095 |
| DP16_RS02840 | Dihydroorotate dehydrogenase | *dtpA* | -2.05901 |
| DP16_RS13710 | DUF47 domain-containing protein | *-* | -2.05462 |
| DP16_RS21040 | HDOD domain-containing protein | *-* | -2.0247 |
| DP16_RS09945 | Mg(2+) transport ATPase C | *mgtC* | -2.02467 |
